# Eocene origin, Miocene diversification and intercontinental dispersal of the genus *Drosera* (Droseraceae)

**DOI:** 10.1101/2020.08.06.240234

**Authors:** Sandeep Sen, Neha Tiwari, R Ganesan

## Abstract

Resolving the evolutionary history of plant carnivory is of great interest to biologists throughout the world. Among the carnivorous plants, Genus *Drosera* (Droseraceae) is highly diverse with a wide pantropical distribution. Despite being a group of interest for evolutionary biology studies since the time of Charles Darwin, the historical biogeography of this group remains poorly understood. In this study, with an improved species sampling from Genbank, we present a reanalyzed phylogenetic hypothesis of the genus *Drosera*. We developed a dated molecular phylogeny of *Drosera* from DNA sequences of nuclear ITS and chloroplast rbcL genes. Divergence times were estimated on the combined dataset using an uncorrelated lognormal relaxed clock model and a known fossil calibration implemented in BEAST. The maximum clade credibility tree was then used for ancestral range estimations using DEC+J model implemented in BioGeoBEARS. Our analysis suggests that *Drosera* evolved during the Mid Eocene 36 Ma [95% HPD: 49.5-26] and have diversified and dispersed from the late Miocene onwards. Ancestral areas estimated using the DEC+J models suggest an African origin followed major radiation within Australia. Diversification in *Drosera* is temporally congruent with the prevailing drier conditions during the Miocene. From Miocene, grasslands and open habitats dominated across continents and might have provided ecological opportunities for their dispersal and diversification. Several long-distance dispersals and range extensions and in situ radiations coinciding with the evolution of drier conditions can explain their extant distribution across continents. Overall our data set provides fresh insights into the biogeographic factors that shaped the origin and evolution of the genus *Drosera*.

## 1. Introduction

The evolution of carnivory in the plant kingdom poses an exciting biological phenomenon and has fascinated biologists ever since Charles Darwin (Briggs 2009, Thorogood 2017). Carnivory has evolved ten times independently in five different orders among the flowering plants (Fleischmann et al., 2018). Carnivorous plants are distributed in open, moist sites, where they acquire the nutrients from trapping insects (Givnish 2014). The remarkable morphological adaptation to trap and digest animal/insect prey poses a selective advantage over other plant lineages to establish them in infertile soils. Understanding of their ancient dispersal pathways and the rate of interchange between different continents in the past is of utmost importance in giving vital clues to the evolution of the species and shed light on the development of its habitat conditions across time and space.

Consisting of ~ 250 species, *Drosera* L. (Droseraceae) is among the speciose carnivorous plant genera with a cosmopolitan distribution and is highly diverse in Australia (Gonella et al., 2016; Rivadavia et al., 2003). *Drosera*, commonly known as “sundews,” exhibits unmatched adaptations to attract, trap, and digest insects for their nutritional requirements (Bothe and Drake 2007). A recent study by Palfalvi et al., (2020) suggests the carnivory within the family Droseraceae evolved as a result of an early genome duplication event. The presence of specialized colorful tentacles in its leaves tipped with nectar glands (flypaper traps), adhesives, and digestive enzymes are used to attract, trap and digest insect prey (Hatcher et al., 2020). The evolution of these flypaper traps is intriguing and might have evolved from a leaf only with adhesive glands after the evolution of nastic and trophic glands (Rivadavia et al., 2012). The sister genera Aldrovanda L. and Dionaea Sol. *ex* J. Ellis share exclusively another type of trapping mechanism known as snap traps (Givnish 2014). Aldrovanda is a floating aquatic species distributed in the old world and Australia, and Dionaea is an endemic genus and adapted to USA’s marshy lands.

The systematics of *Drosera* by Rivadavia et al., (2003) described eleven sections and three subgenera within the genus. Their phylogenetic hypothesis further suggests that genus *Drosera* originated in Africa or Australia, followed by several long-distance dispersals from Australia to Neotropics and the Nearctic (Rivadavia et al., 2003). Besides, the Australian species might have expanded their distribution to Neotropics and Africa. The recent phylogenetic hypothesis of *Drosera* developed by Biswal et al. (2017) discussed various biogeographic scenarios. Their study aimed at providing a better understanding of the existing biogeographic hypotheses for *Drosera* as well as phylogenetic positions of the three species found in India. However, their divergence date estimation and tree topologies lacked resolution, which potentially makes the biogeographic inferences hard to interpret. Thus, a comprehensive biogeographic analysis based on an improved dated phylogenetic hypothesis can provide vital clues in the origin and evolution of this interesting group across major biomes.

The fossil records for insectivorous plants are scarce, except for the genus *Aldrovanda* and family Roridulaceae (Sadowski et al., 2014). For *Droseraceae*, the prime being an early Eocene microfossil aged between 55-38 Ma and another pollen fossil record of *Drosera* appeared is sediments of Miocene aged between 22-5 Ma. (Degereef et al., 1989). However, previous dated phylogenetic hypotheses on *Drosera* using these fossil ages and other calibration schemes produced contrasting age estimates. For example, fossil-based estimates by Magallón et al., (2015) suggested a late cretaceous origin 76.8 Ma [95% HPD: 93-53] of *Drosera*. Later, the likelihood-based time tree developed by Biswal et al. (2017) using the two known fossil dates estimated the age of *Drosera* to be 82 Ma. Similarly, the angiosperm phylogenetic tree developed by Smith and Brown (2018), indicated that the age of *Drosera* to be 63 Ma. Together, these studies suggest an age distribution of *Drosera* from Cretaceous to Oligocene. In addition to the variations in divergence dates proposed by previous studies, an influential study on angiosperm origin by Magallón and Sanderson (2001) suggested that several cosmopolitan angiosperm lineages radiated during recent times but not in the Cretaceous.

In this context, with a lack of consensus among the previous studies, we re-investigated the biogeographic history of genus *Drosera* with a more comprehensive sampling of the nuclear ITS and chloroplast *rbcL* sequences available from the Genbank compared to previous studies. The primary goals of the current study are i) to generate an improved dated phylogenetic hypothesis using three previously defined calibrations and compare them after calculating their marginal likelihoods (MLEs) following a path sampling or stepping stone method. And ii) infer the biogeographical history of *Drosera* and the mechanisms that shaped the origin and spread of the genus across continents. Owing to its association with habitats with drier and nutrient-poor conditions (Brewer and Schlauer 2018), iii) we specifically aim to test whether global aridification from Miocene and Plio-Pleistocene have played a significant role in its diversification and establishment in different biomes?

## 2 Materials and methods

### 2.1 Phylogenetic analysis

The DNA sequences for nuclear ITS and chloroplast *rbcL* for 86 species of *Drosera* and two species of *Nepenthes* were downloaded from the GenBank and then aligned with MAFFTv 7.213 (Katoh and Standley 2013) using L-INS-I (accurate) settings. Sequence gaps in the alignment were treated as missing data. The best fit models and optimum partition schemes were evaluated using the Bayesian information criterion (BIC) and a greedy algorithm employed in PartitionFinder v1.1 (Lanfear et al., 2012). Phylogenetic trees were reconstructed using maximum likelihood (ML) and Bayesian inference (BI). ML tree topologies were developed after 1000 bootstrap replicates in RAXML-HPC V.8 (Stamatakis 2006). A Bayesian inference (BI) was further performed in the aligned data set using the program Mr. Bayesv3.2.6 (Ronquist et al., 2012) implemented on CIPRES Science gateway (Miller et al. 2010). Two independent runs of MCMC generations were run for 50 million generations of Markov Chain Monte Carlo (MCMC) by sampling every 1000 generations until the average standard deviation of split frequencies falls less than 0.01. MCMC chain convergence was further assessed by checking the Estimated sample size (ESS) values. The initial 20% trees from the Mr. Bayes analysis was discarded as burn-in. All the phylogenetic trees were rooted using two outgroups from the genus *Nepenthes*.

### 2.2 Divergence time estimation

For divergence time estimations, we used the program BEAST V 1.8 (Drummond et al., 2012) implemented in CIPRES science gateway V3.3 (Miller et al., 2010). Molecular dates were estimated using the combined dataset of ITS and *rbcL*. The dataset was partitioned by gene, and an uncorrelated lognormal relaxed clock model was employed after unlinking the substitution and clock models and linking trees in BEAUTI.v.18.3 (part of the BEAST package). We tested three different calibration schemes to infer the node ages of the *Drosera* phylogeny. In calibration scheme 1 (C1) the crown age of Droseraceae was constrained between 53-92 Ma with mean age of 76 Ma using a normally distributed age prior since these ranges were estimated previously by Magallón et al., (2015). In calibration scheme 2 (C2), we used the known Dorseraceae pollen records from Eocene to constrain the crown age of Droseraceae (Degreef 1997). A uniform prior was set between 34-56 Ma, which is mentioned in Smith et al. (2017). And in calibration scheme 3 (C3), we place a log-normal prior offset of 63 Ma and a standard deviation equal to 1. This age constraint was derived from the larger seed plant phylogeny developed by Smith and Brown (2018). A starting tree satisfying the age priors was set for all the analyses and were generated using ‘*chronopl*’ function in R using the package ape (Paradis et al., 2004). We removed the subtree slide, Wilson balding, narrow, and wide exchange operators from the XML file to prevent BEAST from exploring the topology space and only to estimate the branch lengths. These three models were then compared using the marginal likelihood estimates following a path sampling/stepping stone method implemented in BEAST. Two independent MCMC runs were performed for 100 million generations by sampling every 1000 steps. Outputs of independent runs for the best model were combined using the software Logcombiner 1.8.2 (Rambaut and Drummond 2010a). The software program TRACER (Rambaut and Drummond 2010b) was used to inspect the estimated sample size (ESS) values and proper mixing. The maximum clade credibility (MCC) tree was generated using tree annotator after discarding 20% of the initial runs are burn in.

### 2.3 Biogeographical Inference

The biogeographic analysis was performed in the R package BioGeoBEARS (Matzke, 2013) in the BEAST MCC tree. To estimate the ancestral ranges at each node, we used the dispersal extinction and cladogenesis (DEC) model and its variant DEC+ J model (Matzke, 2014), which accounts for founder event speciation. Furthermore, the other models implemented in BioGeoBEARS viz. DIVALIKE, BAYAREA, and its J variants were also used to estimate the ancestral ranges. However, this approach received several conceptual criticisms by Ree and Sanmartin (2018), especially on the direct model comparisons between these models based on their likelihood scores. For the biogeographic analysis, we designated seven areas for the ancestral range estimations based on the distribution of *Drosera* i.e Afro-Madagascar (A), Neotropics (B), Nearctic (C), Western Palearctic (D), Eastern Palearctic (E), Southeast Asia (F) and Australasia (G) following the extant distribution of *Drosera* taken from published records and other online resources (Table S1).

## 3. Results

### 3.1 Phylogenetic analysis

Our improved phylogenetic hypothesis based on the combined analysis of chloroplast *rbcL* and nuclear ITS represents the genetic relationships of 85 species of *Drosera*. The combined data sets consist of 2004 bp, of which 1227bp was chloroplast *rbcL*, and 777 bp belong to the nuclear ITS. The models of sequence evolution identified by PartitionFinder varied between all three analytical schemes (Table 1). Genus *Drosera* forms a monophyletic clade sister to *Aldrovanda* and *Dionaea* in BI and ML analysis, as in previous studies (Rivadavia et al. 2003, Magallón et al. 2015). The analysis also recovered two major clades within *Drosera*. Clade 1 consisted of species from Africa, Europe, and Asia (ML bootstrap = 97, posterior probability support=1, (supplementary file Figure S1). Clade 2 consists majorly of members from South Pacific (Australia and New Zealand) and the ancestral species (*D. regia* Stephans) from Africa. We found uncertainties in the placement of the previously defined taxonomic sections, and some taxonomic relationships contradict the previous relationships. This difference can be attributed to larger sequence sampling used in this study compared to the previous works by Rivadavia et al. (2003; 2012). Hence, we refrain from making inferences of the taxonomic sections in this study due to our lack of access to several voucher specimens and leave this section open for interpretation.

**Table 1:** Partition schemes and substitution models used for the Maximum likelihood (ML), Bayesian Inference (BI) and Molecular dating (BEAST)

| Sl.no | Gene | RAXML | Mr. Bayes | BEAST |
| --- | --- | --- | --- | --- |
| 1 | *rbcl* codon position1 | GTR+ G | JC+I+G | TRNef+I+G |
| 2 | *rbcl* codon position2 | GTR+ G | JC+I+G | TRNef+I+G |
| 3 | *rbcl* codon position3 | GTR+ G | GTR+G+I | GTR+I+G |
| 4 | Nuclear ITS | GTR+ G | GTR+G | GTR+G |

### 3.2 Divergence time estimation and biogeographic analysis

The highest likelihood corresponded to the C2 (−19578.03), but this was not significantly different than the marginal likelihood scored for C1 (Table 2). The ESS values ad convergence was better in C2; hence we used these settings for a two replicate MCMC search (see supplementary file Figure S2 for alternative tree topologies). Molecular dating analysis inferred that *Droseraceae* diverged from its sister group during Eocene between 42 Ma [95% HPD: 34-54 Ma] (Figure 1). The crown age estimate of genus *Drosera* was 36Ma [95% HPD: 26-49 Ma]. Split between the two main clades within *Drosera* occurred during the Oligocene 30Ma [95% HPD: 21-42 Ma]. DEC+J model was favored amongst all other models (-lnl=130.069, AICc= 266.4; Table 3) and the null model rejected at p< 0.0001. All the ancestral range estimations identified an African origin of *Drosera* and a further dispersal to Australia during the Eocene (Node 1; Figures 1, 2; Table 4). Further, the genus *Drosera* had undergone massive radiation within Australia, starting from Oligocene. These radiations were followed by several dispersals to Neotropics and Africa starting from the Mid Miocene (Node numbers 2, 3, 4, 5, 6; Figures 1, 2). A single back dispersal to Africa and dispersal to Southeast Asia from Australia was observed during this period (Node numbers 7, 8, 9; Figures 1, 2). Furthermore, we observed several *in situ* radiations within the Neotropics, and the Afro-Madagascar in the late Miocene, followed by dispersal from Australia (Node numbers 10,11,12; Figures 1, 2).

**Figure 1:**
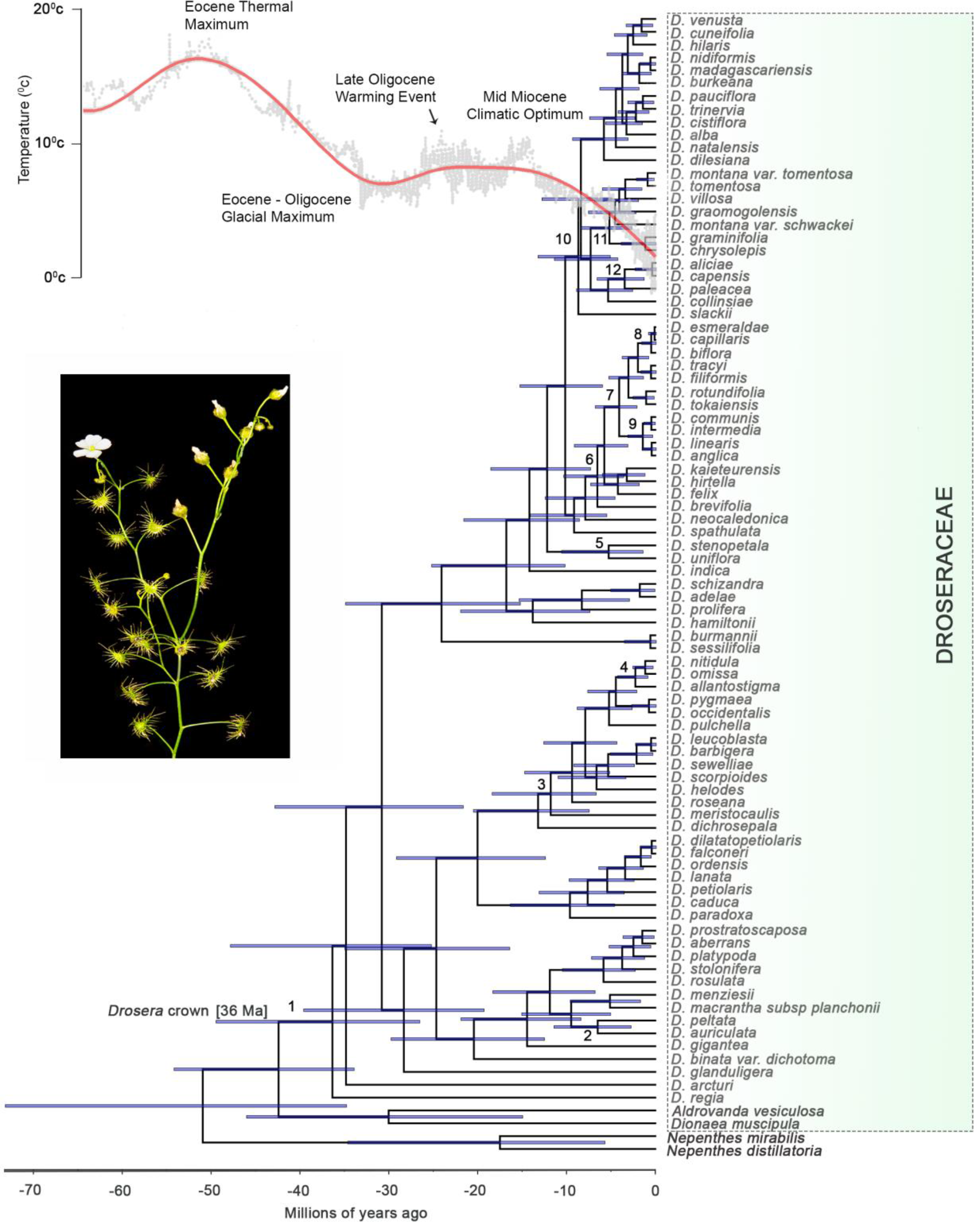
Maximum clade credibility (MCC) chronogram from molecular dating analysis of *Drosera*. The temperature curve represents the major trends in climate from Cenozoic (−65Ma) to present (0Ma) which is derived from oxygen isotope (Δ_18_O) data from Zachos et al., (2008) were converted to absolute temperatures. The divergence time and HPD intervals of the numbered nodes are represented in Table 4. Inset: *D. indica*.

**Figure 2:**
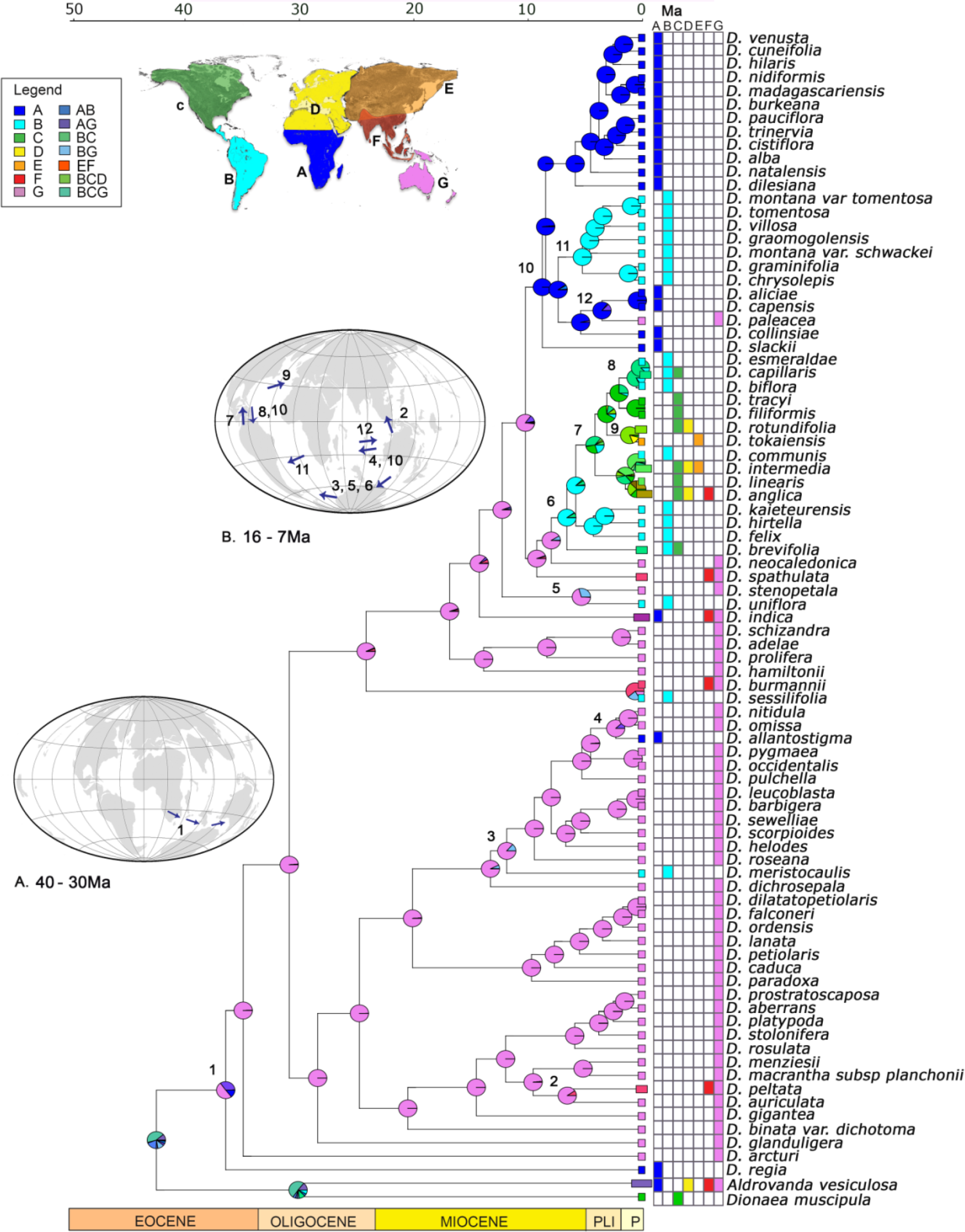
Historical biogeography of *Drosera*. Colored boxes to the left of species names shows current geographical distributions indicated in the distribution map. Pie charts on the nodes represents probabilities of the most likely ancestral ranges estimated from the DEC+J model. The relative continental positions are represented in the globes and the arrows and the numbers represents important dispersal events.

**Table 2:** BEAST model comparisons based on three different calibration schemes used in this study

|  | C1 | C2 | C3 |
| --- | --- | --- | --- |
| Path sampling | -19676.385 | -19578.03 | -19594.457 |
| Stepping stone | -19671.036 | -19486.71 | -19542.725 |

**Table 3:** Model comparisons in BioGeoBEARS.

| Model | -LNL | No. of parameters | *d* | *e* | *j* | *AICc* | *p* |
| --- | --- | --- | --- | --- | --- | --- | --- |
| DEC | -141.26 | 2 | 0.016 | 0.002 | 0 | 286.7 |  |
| **DEC+j** | **-130.069** | **3** | **0.010** | **1X10<sup>-12</sup>** | **0.032** | **266.4** | **>0.0001** |
| DIVALIKE | -138.348 | 2 | 0.018 | 1X10<sup>-12</sup> | 0 | 280.8 |  |
| DIVALIKE+j | -132.404 | 3 | 0.011 | 1X10<sup>-12</sup> | 0.029 | 271.1 | 0.0006 |
| BAYAREA | -182.473 | 2 | 0.016 | 0.043 | 0 | 369.1 |  |
| BAYAREA+j | -136.858 | 3 | 0.008 | 0.001 | 0.046 | 280 | >0.0001 |

d=rate of dispersal; e= rate of extinction; j= relative probability of founder event speciation

**Table 4:** Divergence time and the distribution of its highest posterior density (HPD) intervals of biogeographically important nodes. Node numbers correspond to the ones in Figures 1, 2, 2A, and 2B.

| Node | Description | Divergence time (Ma) |  |
| --- | --- | --- | --- |
|  |  | Mean | 95%HPD |
| 1 | *Drosera* crown | 36.4 | 49.5 - 26 |
| 2 | Split: *D. peltata* and *D. auriculata* | 6.5 | 11.5 – 2.8 |
| 3 | Stem of *D. meristocaulis* | 11.8 | 18.4 – 6.7 |
| 4 | Stem of *D. allantostigma* | 2.3 | 4.3 - 0.9 |
| 5 | Split: *D. stenopetala* and<br>*D. uniflora* | 5.3 | 10.6 – 1.4 |
| 6 | Stem *D. brevifolia* | 6.6 | 10.4 -3.6 |
| 7 | Crown of the Nearctic clade | 4.1 | 6.8 – 2.7 |
| 8 | Stem of *D. biflora* | 0.5 | 1.6 - 0.2 |
| 9 | Split: *D. tokaiensis* – *D. rotundifolia* | 1.1 | 2.6 -0.2 |
| 10 | Crown of the clade dispersed to Africa<br>from Australia | 8.7 | 13.2 – 5.1 |
| 11 | Stem of the clade dispersed to<br>Neotropics from Africa | 5.2 | 8.4 – 2.7 |
| 12 | Stem of *D. paleacea* | 3.5 | 6.6 – 1.3 |

## 4. Discussion

Our results support a complex biogeographic history of genus *Drosera* and explain the historical factors that influenced its dispersal and diversification patterns. It consists of an African origin followed by Australian radiations, long-distance dispersals, range extensions, and several *in situ* radiations in other continents. *In situ* radiations in multiple lineages were followed by the intensification of grassland spread and reductions in global temperatures. The implications of our new, improved analysis and its potential shortcomings are discussed herein.

### 4.1. Origin and historical spread of *Drosera*

The results from our time-calibrated phylogeny suggest that the ancestors of *Drosera* have evolved in Africa during the Eocene and dispersed to Australia as hypothesized by Rivadavia et al., (2003). The molecular clock dated this event to Eocene, and the time corresponds well to the Eocene-Oligocene glacial maximum. This period exhibited arid conditions in the southern regions of Australia and Africa (Buerki et al., 2013), which might have provided a suitable environment for *Drosera* to disperse from Africa to Australia via Antarctica which was once a common route for several lineages to disperse into Australia (Benda et al., 2019). For example, Sapindaceous lineages were shown to have using this dispersal route from Australia to Africa during the Middle Paleocene-Late Eocene (Buerki et al., 2013). These age estimates are too young to invoke a Gondwanan vicariance hypothesis. And another possibility is dispersal through the now-submerged Crozet plateaus and Kerguelen islands to Australia (see Kayaalp et al., 2017, Renner et al., 2010).

The dispersals from Africa were followed by tremendous in situ radiations in *Drosera* within the Australian continent. Starting from the Oligocene, a monsoonal drier condition has evolved in northern Australia. Due to the drop in global temperature, there was a reduction in the extent of broad-leaved forests. Previous studies have shown that these changes are known to affect the native vegetation of Australia. For example, in situ radiations in Eucalyptus subgenus Symphyomyrtus (Ladiges et al., 2003) coincides well with this time frame. The early to mid-Miocene weather was mostly warm, followed by cool and dry conditions in the late Miocene. Especially the fossil pollen from freshwater swamps suggests the existence of a relatively open heath scrubland, which has highly acidic infertile soils with the presence of repeated wildfire. The late Miocene witnessed a reduction in rainforests and an increase in Eucalyptus forests, indicating a colder and drier condition. And the existing rainforests continued to reduce with an increase in grassland and arid climate in Pliocene (Martin 2006). Our phylogenetic hypotheses suggesting several species radiations in Australia is highly congruent with this period. As shown elsewhere (Givnish 2014), these unfavorable conditions are advantageous for a carnivorous plant genus like *Drosera*. Moreover, during this time frame, a change in climate to drier state and the incidence of fire and the spread of open grasslands might have provided several new niches for *Drosera* to diversify within the Australian continent (see Lamont and Downes 2011).

The diversification in *Drosera* also corresponds well to the variation in global climatic trends derived from Δ_18_O data to the Cenozoic to present (Zachos et al., 2008), which was evident from Oligocene-Miocene radiations in Australia (Figure 1). These radiations in Australia were followed by several long-distance dispersals from Australia to South America and range extensions to Southeast Asia (Node numbers 2, 3, 4, and 5). For example, *D. meristocaulis* Maguire & Wurdack, is shown to have undergone dispersal from Australia to the Neotropics in the Mid-Miocene. Our biogeographic analysis and molecular dates are also in agreement with a possible long-distance dispersal from Australia to Neotropics (see Thornhill et al., 2015). These dispersals are intriguing, but, can take place via Antarctica (35-10Ma) and possibly mediated through an establishment of the West Wind Drift (WWD) and equatorial currents during this period (see Buerki et al., 2013). And this long-distance dispersal can also be through the islands that have existed in the Pacific basin since the Oligocene, such as Hawaiian archipelago and Fiji Islands (Chen et al., 2014). However, such long-distance dispersal from Australia to Neotropics via Antarctica is not entirely rare. Similar patterns are found in the *Ampelopsis* clade of Vitaceae (Nie et al. 2012) and the cosmopolitan genus *Ranunculus* (Emadzade et al., 2011). To better understand the major dispersal events, we marked the results in Figure 2 (insets A and B).

A lineage dispersed to Neotropics from Australia (Node 6), has undergone *in situ* radiation and then dispersed to Nearctic and Western Palearctic (Node 7, 9) during the Miocene. A back dispersal then followed this dispersal to Neotropics (Node 8). Interestingly, our age estimates of these nodes align well will the expansion of grassland biomes within the Neotropics and the Nearctic (see Fig 6c in Meseguer et al., 2015). Further, there were long-distance dispersals from Australia to Africa during the Miocene (node 10) and might be a result of via wind or ocean currents. (Dick et al., 2007). During the late Miocene, Africa also several ecological transitions, mainly the spread of C4 type grasslands; overall, these transitions might have been favorable for *Drosera* to diversify (Meseguer et al., 2015).

We observe a single dispersal from Africa to South America during the Miocene, followed by species radiation in that lineage (Node 12). This pattern of *in situ* radiation again corresponds well with the timing of grassland expansion in South America, which could be a favorable time for *Drosera* to diversify within. *Drosera paleacea* (Node 12) originate via dispersal from Africa to Australia during the late Miocene. A long-distance dispersal from Africa to Australia remains the most likely explanation supported by well-studied taxa such as Simaroubaceae (Clayton et al., 2009). Overall findings from this study suggest the role of then prevailing drier conditions from Miocene and long-distance dispersals (possibly by birds) explains the inter-continental dispersals and radiations in the genus *Drosera*.

### 4.2 Molecular dating and previous hypotheses

Maximum likelihood RelTime time estimates of *Drosera* by Biswal et al., (2017) computed based on branch lengths optimized by ML (Tamura et al., 2012) lack resolution and was also characterized by several polytomies. Usage of the second fossil calibration (25-5Ma) was also not well justified. and we refrained from adding it in our divergence dates due to the confusion in its placement. The RelTime method is computationally efficient for estimating divergence times (Mellow et al., 2019). Their dated tree topology resulted in several polytomies and raised practical difficulties in interpreting the results (See Fig. 7 Biswal et al., 2017). An incomplete taxon sampling from the GenBank might be another reason that prevented the authors from developing a resolved tree topology.

Further, inadequate taxon sampling is also known to underestimate the divergence times (Linder et al., 2005). A densely sampled phylogeny and genome sequences can improve the taxonomic relationships and biogeographic inferences within the genus *Drosera*, which is a way forward. Interestingly, our molecular dating approaches suggest an Eocene origin of the family *Drosera*, opposing the previous date estimates. Besides, our newly inferred dates are more congruent with the global decrease in temperature, which might have set the stage for diversification of *Drosera* in different biomes. It is also noteworthy that the recent large scale dated phylogeny developed by Ramírez-Barahona et al., (2020) also identified a Cretaceous origin of Droseraceae 65.8 M instead of an Eocene origin as per our dating analysis, but with a larger credible interval spanning from 30-94 Ma. We did not include these in our calibration schemes, as these ages ranges were covered in C1 and C3.

## 5. Conclusions

Aridifications can exert intense selective pressure, influencing diversification in plant lineages (Gutiérrez-Ortega et al., 2017). Our analysis confirms an Eocene origin and Miocene diversification of *Drosera* followed by a complex dispersal pathway, including long-distance dispersals, to establish its presence across continents. *Drosera*’s diversification is temporally congruent with the aridification and presence of open grassland and fire in Australia, Africa, and Neotropics. Our study also confirms that *Drosera* exhibits an early history of range expansion between major continental areas. Overall, our finding agrees with previously discussed dispersal pathways for major angiosperm lineages (Buerki et al., 2013; Chen et al., 2014). Our results will have implications for understanding the biogeographic history of *Drosera* and also expected to shed light on the evolution of modern-day biomes. Further studies with a more extensive taxon sampling and large sequence datasets are required to deepen the knowledge in phylogeny, biogeographic history and diversification dynamics of *Drosera*.

## Acknowledgements

Research grants from the Department of Biotechnology, India for the research Project Bioresource and Sustainable Development in Northeast India (BT/01/17/NE/TAX Dt 29 March 2018), and the Biodiversity exploration grant from Christopher Davidson (Flora of the World) is acknowledged. We thank Divya, Siddarthan Surveswaran, and Priyadarsanan for their comments. We thank Abu Hang Samuel for sharing his field photographs with us.

