## Supplemental file for "Eocene origin, Miocene diversification and intercontinental dispersal of the genus *Drosera* (Droseraceae)"

**Table S1**: List of taxa used in this study and the corresponding accession numbers from GenBank. The species ranges are denoted as A= Afro-Madagascar, B= Neotropics, C= Nearctic, D= Western palearctic, E= Eastern Palearctic, F= Southeast Asia and G= Australasia

| **Taxon** | **NCBI Accession Number** | | **Species Range** |
|---|---|---|---|
| | ITS | rbcL | |
| *D. aberrans* (Lowrie & Carlquist) Lowrie & Conran | KP171689.1 | KP268949.1 | G |
| *D. adelae* F.Muell. | | AY096107.1 | G |
| *D. alba* Phillips | | AB072515.1 | A |
| *D. aliciae* Raym.-Hamet | | AB072516.1 | A |
| *D. allantostigma* (N.G Marchant & Lowrie) Lowrie & Conran | KP171687.1 | KP268945.1 | A |
| *D. anglica* Huds. | AB355666.1 | AB355692.1 | C, D, F |
| *D.* arcturi Hook. | | AB072512.1 | G |
| *D. auriculata* | KP171688.1 | KP268948.1 | G |
| *D. barbigera* Planch. | JQ712490.1 | JQ712489.1 | G |
| *D. biflora* Willd. Ex Roem. & Schult. | | AB072518.1 | B |
| *D. binata* Labill. | HM204879.1 | AB072922.1 | G |
| *D. brevifolia* Pursh | JN388055.1 | AB072519.1 | B, C |
| *D. brukeana* Planch | | AB072520.1 | A |
| *D. burmanii* Vahl | JN388056.1 | KT794003 | F, G |
| *D. caduca* Lowrie | | AB072510.1 | G |
| *D. capensis* L. | HM204880.1 | AB917049.1 | A |
| *D. capillaris* Poir. | JN388059.1 | KJ773463.1 | B, C |
| *D. chrysolepis* Taub | JN388060.1 | AB072522.1 | B |
| *D. cisitiflora* L. | | AB072523.1 | A |
| *D. collinsiae* N.E.Br | JN388061.1 | AB072524.1 | A |
| *D. communis* A.St.-Hil. | JN388070.1 | | B |
| *D. cuneifolia* L.f. | | AB072525.1 | A |
| *D. dichrosepala* Turcz. | | L01910.2 | G |
| *D. dilatato-petolaris* K.Kondo | KP171680.1 | KP268954.1 | G |
| *D. dilesiana* Exell & J.R Laundon | JN388062.1 | | A |
| *D. esmeraldae* (Steyerm.) Maguire & Wurdack | | AB072526 | B |
| *D. falconeri* Tsang ex. K.Kondo | HM204882.1 | KP268943.1 | G |
| *D. felix* Steyerm. & L.B.Sm. | | AB072527.1 | B |
| *D. filiformis* Raf. | JN388063.1 | KJ773464.1 | C |
| *D. gigantea* Lindl. | | L19528.2 | G |
| *D. glanduligera* Lehm. | JN388039.1 | AB072511.1 | G |
| *D. graminifolia* A.St-Hill. | JN388064.1 | AB072528.1 | B |
| *D. graomogolensis* T.R.S. Silva | JN388065.1 | AB072529.1 | B |
| *D. hamiltonii* C.R.P. Andrews | HM204884.1 | AB072921.1 | G |
| *D. helodes* N.G Marchant & Lowrie | KP171690.1 | KP268950.1 | G |

| | | | |
|---|---|---|---|
| *D. hirtella* A.St.-Hil. | JN388066.1 | AB072531.1 | B |
| *D. indica* L. | JN388067.1 | L19529.2 | A, F, G |
| *D. intermedia* Hayne | JN388069.1 | JN891175.1 | C, B, D |
| *D. kaieteurensis* Brumm.-Ding. | MF785371.1 | AB072532.1 | B |
| *D. lanata* K.Kondo | KP171685.1 | KP268942.1 | G |
| *D. linearis* Goldie | MG237379.1 | MG246288.1 | C |
| *D. macrantha subsp. Planchonii* Endl. | | AB072549 | G |
| *D. madagascariensis* DC. | JN388071.1 | AB072533.1 | A |
| *D. menziesii* R.Br.ex DC. | KP171679.1 | KP268947.1 | G |
| *D. meristocaulis* Maguire & Wurdack | JN388038.1 | JN388035.1 | B |
| *D. montana* A.St-Hill. | JN388072.1 | AB072536.1 | B |
| *D. muscipula* | JN388078.1 | | |
| *D. natalensis* Diels | JN388073.1 | AB072537.1 | A |
| *D. nidiformis* Debbert | HM204885.1 | | A |
| *D. nitidula* Planch. | JN388040.1 | JN388036.1 | G |
| *D. occidentalis* Morrison | JN388042.1 | AB072506.1 | G |
| *D. omissa* Diels | KP171686.1 | KP268944.1 | G |
| *D. ordensis* Lowrie | JN388075.1 | JN388037.1 | G |
| *D. paleaceae* DC | HM204886.1 | KP268941.1 | G |
| *D. paradoxa* Lowrie | JN388043.1 | | G |
| *D. pauciflora* Banks ex DC | | AB072552.1 | A |
| *D. pelteta* Thunb. | KF016002.1 | KT794002 | F, G |
| *D. petolaris* R.Br. ex DC. | | L01913.2 | G |
| *D. platypoda* Turcz. | | AB072547.1 | G |
| *D. prolifera* C.T.White | KP171678.1 | KP268938.1 | G |
| *D. prostratoscaposa* Lowrie & Carlquist | | AB072554.1 | G |
| *D. pulchella* Lehm. | JN388076.1 | KP268939.1 | G |
| *D. regia* Stephens | JN388077.1 | L01914.2 | A |
| *D. roseana* N.G Marchant & Lowrie | KP171691.1 | KP268952.1 | G |
| *D. rotundifolia* L. | AB355664.1 | AB355691.1 | C, D |
| *D. schizandra* Diels | | KP268953.1 | G |
| *D. scorpioides* Planch. | JN388041.1 | AB072509.1 | G |
| *D. sessilifolia* A.St.-Hil. | JN388057.1 | AB072551.1 | B |
| *D. sewelliae* Diels | | KP268951.1 | G |
| *D. slackii* Cheek | HM204889.1 | | A |
| *D. spathulate* Labill. | AB355671.1 | AB355696.1 | F, G |
| *D. stenopetala* Hook.f. | | AB072539.1 | G |
| *D. tokaiensis* (Komiya & C. Shibata) T.Nakamura & E.Ueda | AB355687.1 | AB355698.1 | E |
| *D. tomentosa* A.St-Hill. | JN388053.1 | | B |
| *D. tracyi* | | KJ773465.1 | C |
| *D. trinervia* Spreng. | | AB072553.1 | A |
| *D. uniflora* Willd. | | AB072540.1 | B |

| | | | |
|---|---|---|---|
| *D. venusta* Debbert | | AB917048.1 | A |
| *D. villosa* A.St-Hill. | JN388054.1 | | B |
| *Dionaea muscipula* J. Ellis | HM204877.1 | MH749062.1 | C |
| *D. hilaris* Cham. & Schltdl. | | AB072530.1 | A |
| *D. leucoblasta* Benth. | | *KP268946* | G |
| *D. occidentalis var. microscapa (Debbert) Schlauer* | | *AB072506* | G |
| *D. neocaledonica* Raym.-Hamet | AB072544.1 | | G |
| *D. pygmaea* DC | | AB072505.1 | G |
| *D. rosulata* Lehm | | AB072555.1 | G |
| *D. schwackei* (Diels) Rivadavia | | AB072535 | B |
| *D. stolonifera* Endl. | | L19531.2 | G |
| **Outgroups** | | | |
| *Nepenthes vieillardii* Hook. | AB769065.1 | AB103323.1 | G |
| *Nepenthes distillatoria Graham* | AB675877.1 | MK059425.1 | F |
| *Aldrovanda vesiculosa* L. | AB332395.1 | AY096106.1 | A,D,F,G |

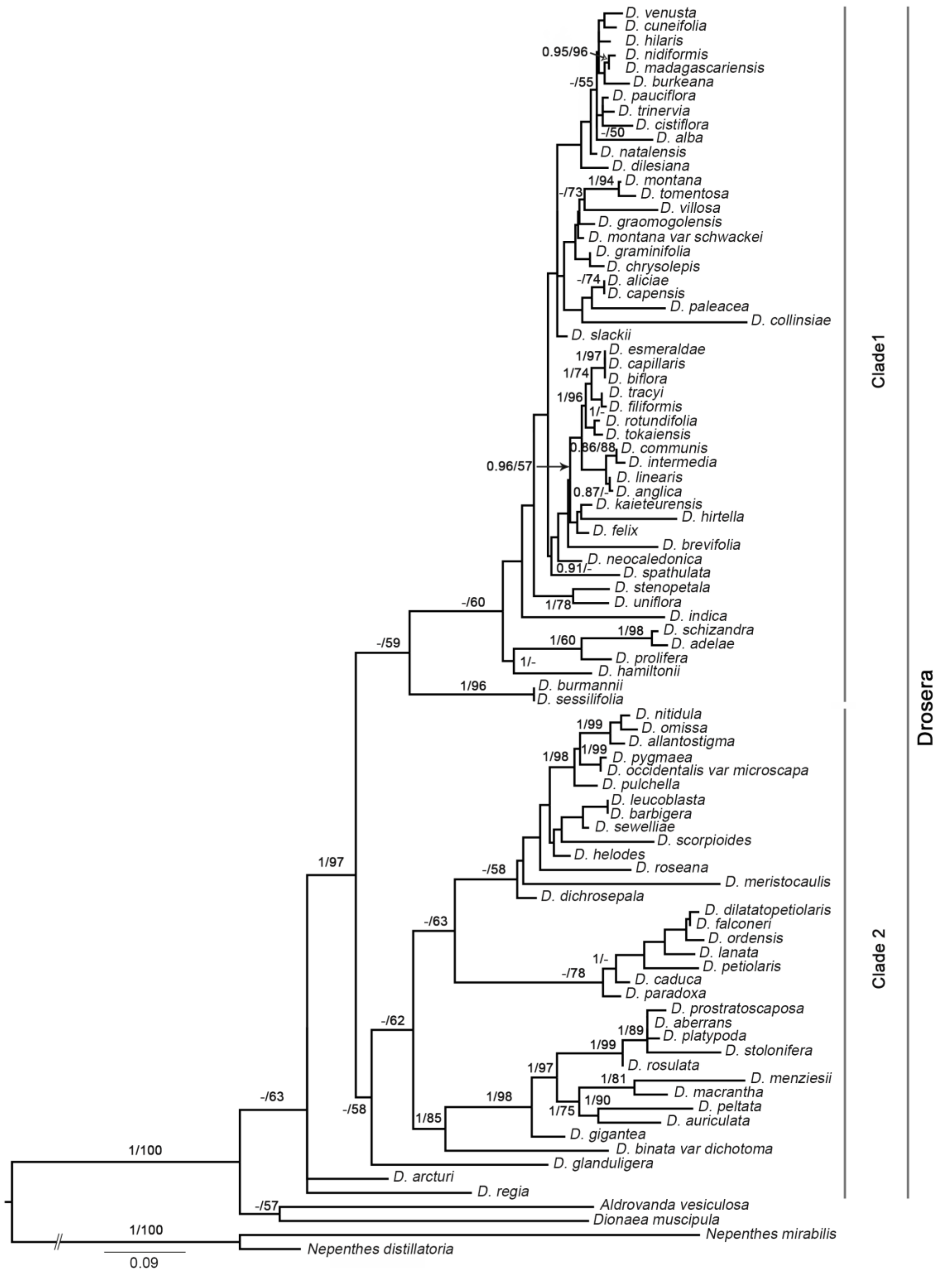

Figure S1: Maximum likelihood phylogenetic tree topology of *Drosera* derived from the concatenated dataset. Support values branches represent Bayesian posterior probabilities (BPP) and maximum likelihood bootstrap values (MLBS). Values below MLBS 50 and BPP 0.95 are not shown.

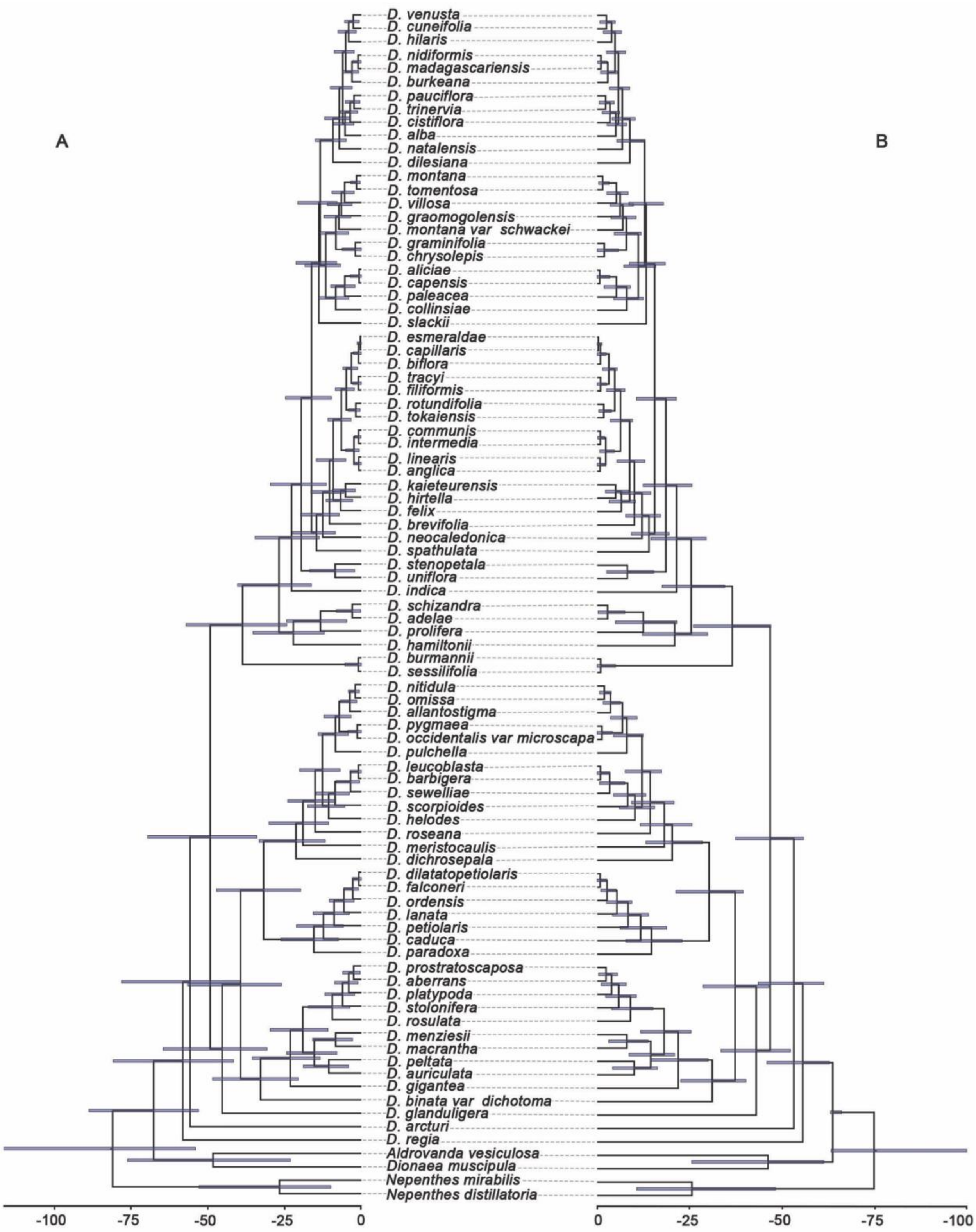

**Figure S2**: BEAST MCC tree obtained from secondary calibration analysis A) from Magallón et al., 2015 (C1) and B) from Smith and Brown 2018 (C3). Blue horizontal bars represent 95% highest posterior density (HPD) intervals
